# Mitochondrial F_1_F_O_ ATP synthase determines the local proton motive force in cristae tips

**DOI:** 10.1101/2021.02.11.430746

**Authors:** Bettina Rieger, Tasnim Arroum, Jimmy Villalta, Karin B. Busch

## Abstract

The classical view of oxidative phosphorylation is that a proton motive force PMF generated by the respiratory chain complexes fuels ATP synthesis. Under glycolytic conditions, ATP synthase in its reverse mode also can contribute to the PMF. Here, we dissected the two functions of ATP synthase and the role of its inhibitory factor 1 (IF1) under different metabolic conditions in detail. pH profiles of mitochondrial sub-compartments were recorded with high spatial resolution in live mammalian cells by positioning a pH-sensor directly at F_1_ and F_O_ of ATP synthase, complex IV and in the matrix. Our results clearly show that ATP synthase activity is substantially controlling the PMF and that IF1 is essential under OXPHOS conditions to prevent reverse ATP synthase activity due to an almost negligible ΔpH.

**GRAPHICAL ABSTRACT:** 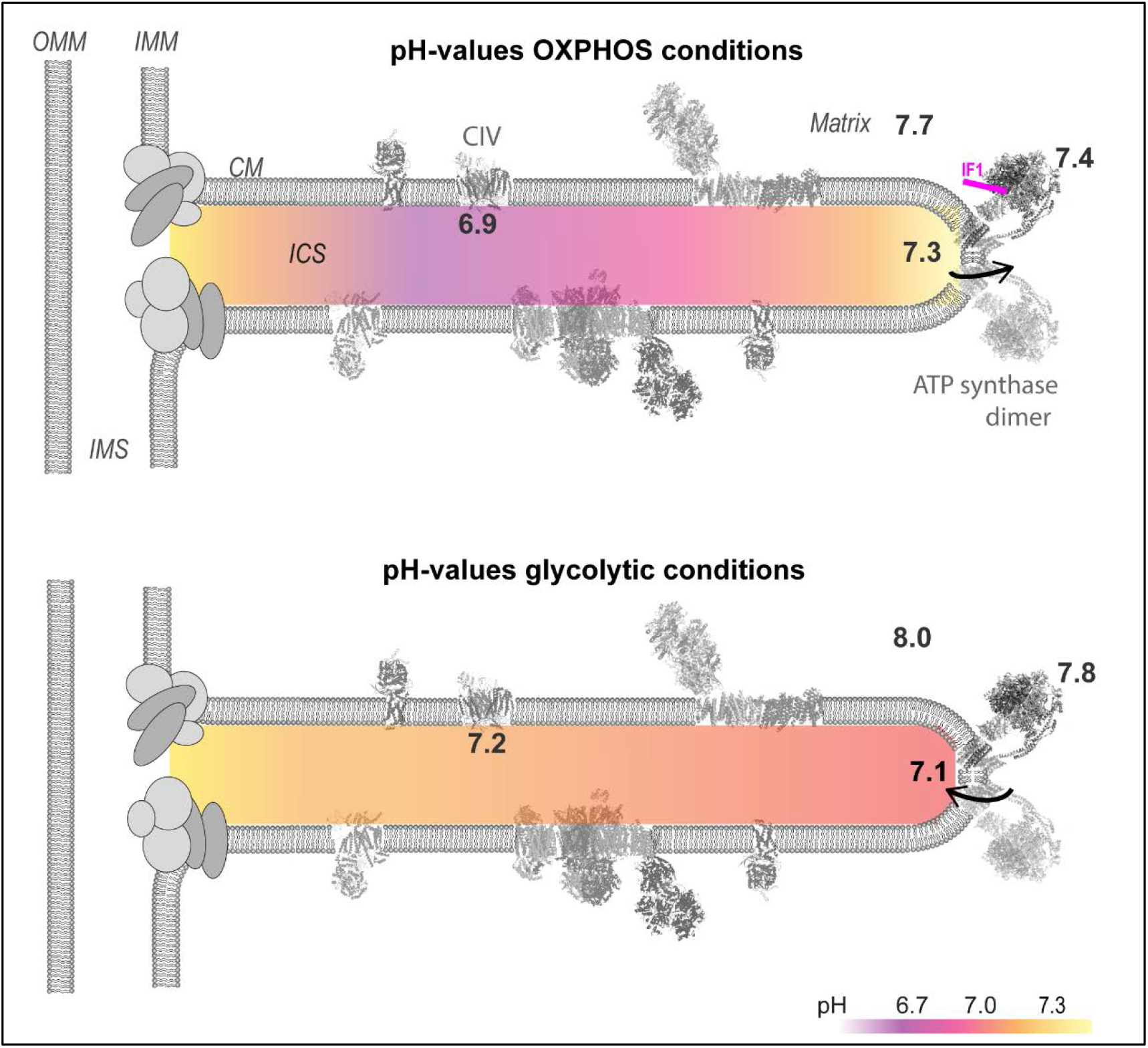

**HIGHLIGHTS:**

- The ΔpH along and across the inner mitochondrial membrane is not homogeneous
- The proton motive force at cristae tips is controlled by F_1_ F_O_ ATP synthase
- Under OXPHOS conditions, the pH difference between F_O_ and F_1_ of active ATP synthase is almost negligible (1.2 proton vs. 1 proton equivalent)
- IF1 is required to prevent the onset of ATP hydrolysis under OXPHOS conditions

## Introduction

Mitochondria are cellular power plants that are the main source of ATP under respiratory conditions. The respiratory complexes CI (NADH-dehydrogenase, NADH: Coenzym Q Oxidoreduktase) and CII (succinate-dehydrogenase) deliver electrons to CIII (cytochrome *c* reductase, coenzyme Q: cytochrome *c* – oxidoreductase). The final electron acceptor of the electron transport chain (ETC) is CIV (cytochrome *c* oxidase). The redox activity of CI, CIII and CIV is coupled to proton pumping from the matrix to the intracristal space (ICS). Thereby, a proton motive force is generated which is the transmembrane difference of electrochemical potential of protons (ΔμH^+^). In his chemiosmotic theory, Peter Mitchell (Mitchell, 1961, Mitchell, 1966, Mitchell & Moyle, 1967) defined the proton motive force Δ*p* (Δ*p*=ΔµH*+*/F, F=Faraday constant), also known as PMF. Δ*p* consists of an electric (ΔΨ_m_) and a chemical part (ΔpH_m_): Δ*p* = ΔΨ_m_ -2.3RT/F ΔpH (mV). Foremost, the Δ*p* drives ATP synthesis, coupling the oxidative part with ADP phosphorylation by F_1_F_O_ ATP synthase in a process known as oxidative phosphorylation (OXPHOS). While the primary proton pumps of the respiratory chain are localized in cristae sheets, rows of dimers of ATP synthase line up along the cristae rims (Blum, Hahn et al., 2019, Davies, Anselmi et al., 2012, Davies, Strauss et al., 2011, Gilkerson, Selker et al., 2003, Vogel, Bornhovd et al., 2006). F_1_F_O_ ATP synthase is constituted of a subcomplex in the membrane, the proton pump, F_O_, and a hydrophilic part extending into the matrix, the F_1_ part. Proton flow from the IMS into the matrix drives ADP phosphorylation (ATP synthase activity). This reaction is reversible and ATP synthase can function as an ATP hydrolase. In order to block futile ATP hydrolysis, e.g. by unassembled F_1_, many cells express the inhibitory factor 1 (IF1) (Campanella, Casswell et al., 2008, Fujikawa, Imamura et al., 2012, Gledhill, Montgomery et al., 2007). IF1 activity is regulated by phosphorylation (Garcia-Bermudez, Sanchez-Arago et al., 2015), Ca^2+^ and protons, making IF1 responsive to key functional parameters inside mitochondria (Boreikaite, Wicky et al., 2019) and thus a master regulator of bioenergetics (Barbato, Sgarbi et al., 2015, Garcia-Bermudez & Cuezva, 2016). The question, whether IF1 modulates ATP synthase in addition to hydrolysis activity *in cellulo* is still under debate (Garcia-Aguilar & Cuezva, 2018, Gledhill et al., 2007). Moreover, the effect of IF1 on the activity of ATP synthase and local Δ*p* has not been resolved so far, as it is technically challenging to determine pH values and pH dynamics in mitochondrial sub-compartments. However, to dissect ATP synthesis and link it to local Δ*p*, this is an essential information.

In order to proceed our knowledge in this matter, we determined local pH values at ATP synthase with high resolution at the *p*- and *n*-side of the enzyme. Local pH determination at opposite sides of the enzyme provides information about the activity of ATP synthase, allows to resolve forward and reverse activity *in situ* and link it to the *Δp*. To do so, a pH-sensitive GFP was introduced to the F_1_ subcomplex and the F_0_ subcomplex via genetic fusion. This allowed to determine pH values in the IMS (*p*-side)and on the matrix side (*n*-side) of the ATP synthase. To understand the role of IF1 in controlling ATP synthase activity, IF1-knockout cells and cells expressing constitutively active IF1-H49K (Garcia-Aguilar & Cuezva, 2018, Schnizer, Van Heeke et al., 1996) were generated. For on-side pH-determination, the pH-sensitive GFP derivative sEcGFP (Orij, Postmus et al., 2009), also known as pHLuorin, was used as a ratio-metric pH sensor (Gao, Knight et al., 2004, Rieger, Junge et al., 2014). The generated pH profiles revealed that the local Δ*p* was unexpectedly low under OXPHOS, that is ATP synthesis conditions. Moreover, our data clearly show that IF1 is necessary to block ATP hydrolysis under OXPHOS conditions. Since the local Δ*p* at active ATP synthase is low under steady state OXPHOS, the initiation of reverse ATPase mode must be prevented. In accordance with this, the Δ*p* was maximal in respiring cells with a constitutively active IF1 variant (IF1-H49K).

## RESULTS

HeLa cells were chosen as a suitable model to study low and high mitochondrial OXPHOS activity. A switch is achieved by changing the carbohydrate supply (Rossignol, Gilkerson et al., 2004). Comparison of the metabolic profile of cells grown with high glucose (HGlc: 25 mM) and galactose (10 mM) for 3 weeks increased respiration on cost of glycolysis as determined with an automatic flux analyzer that records oxygen consumption rates (OCR) as proxy for OXPHOS activity and extracellular acidification rates (ECAR) as proxy for glycolytic activity. Cells grown with glucose were characterized by a low OCR/ECAR ratio, while cells grown with galactose containing medium had a high OCR/ECAR ratio (Supplementary Figure S1a). Following, cells with high OCR/ECAR ratio are also called OXPHOS cells.

### pH measurements directly at ATP synthase allow the discrimination of ATP synthase and ATPase activity in situ

To dissect, how ATP synthase activity feeds back to the PMF, we placed local pH sensors and recorded intramitochondrial pH values. As pH sensor, the pH-sensitive GFP derivative sEcGFP (pHLuorin, superecliptic GFP) (Miesenbock, De Angelis et al., 1998) was used. Superecliptic GFP is a pH sensitive variant from the *Aquorea victoria* green fluorescent protein, with 9 mutations that lead to increased fluorescence and ratio metric pH sensitivity in comparison to the original ecliptic variant. Since the pH-dependent emission spectra display an isosbestic point, sEcGFP can be used as a ratio metric pH sensor(Gao et al., 2004, Rieger et al., 2014) (Figure 1A). Different site-specific pH sensors were generated by genetically fusing sEcGFP to subunits of CV, CIV and targeting it to the matrix, respectively (Figure 1B). sEcGFP was fused to subunit γ (SU γ) of the F_1_ subcomplex of CV at the *n*-side of the IMM, to subunit e (SU e) of F_O_ subcomplex of CV at the *p*-side of the cristae membrane and to subunit CoxVIIIa of CIV at the *p*-side of the IMM(Rosselin, Santo-Domingo et al., 2017). The correct localizations have been confirmed by immuno-EM in a different study (Rosselin et al., 2017). After excitation with 405 nm, the emission in two channels was simultaneously recorded and the ratio calculated (λ_511_/λ_464_). First, pH calibration curves were generated by incubation of cells in media with different pH values. Addition of ATP synthase inhibitor oligomycin and uncouplers (Carbonylcyanid-p-trifluoromethoxyphenylhydrazon =FCCP, Nigericin) resulted in the equilibration of mitochondrial with external pH (Figure 1C). Detergents were not required for pH equilibration (supplementary Figure S2). The response of cells expressing mt-sEcGFP, SU γ-sEcGFP and SU e-sEcGFP to different pH values were similar (Figure 1D). The color-coded ratio metric images of mt-sEcGFP expressing cells show the response to pH changes in a false color depiction (Figure1D, right panel).To test the feasibility for recording physiological pH under different metabolic conditions, the pH at SU e, at SU γ and in the matrix was determined in glycolytic and respiratory cells. Under glycolytic conditions, the pH at SU e was pH = 7.06 ± 0.25 (Figure 1E). At CIV subunit CoxVIIIa, the pH was significantly higher (pH=7.21 ± 0.26). Based on the small OCR/ECAR, we explain the lower pH at SU e by reverse ATP synthase (ATPase) activity. ATP hydrolysis fuels pumping of protons from the matrix into the intracristal space. In the matrix, the pH at SU γ of CV and in the bulk were both alkaline (pH_matrix_ = 7.99 ± 0.09) (Figure 1G). Addition of oligomycin and uncouplers increased the pH in the ICS and decreased the pH in the matrix, finally reaching the pH of the extracellular medium (Figure 1E,F). The proton concentration at SU e was higher under respiratory conditions (pH = 7.28 ± 0.24) compared to glycolytic conditions (Figure 1G,H). Surprisingly, the pH at SU γ and in the matrix were more acidic under OXPHOS conditions in line with proton pumping into the matrix by ATP synthase. At CIV, the pH was significantly higher under glycolytic conditions (pH=7.21 ± 0.26) than under respiratory conditions (pH=6.88 ± 0.15, *p*=8.6^-11^) due to increased proton pumping activity of CIV under respiratory (OXPHOS) conditions (Figure 1I).

**Figure 1.**
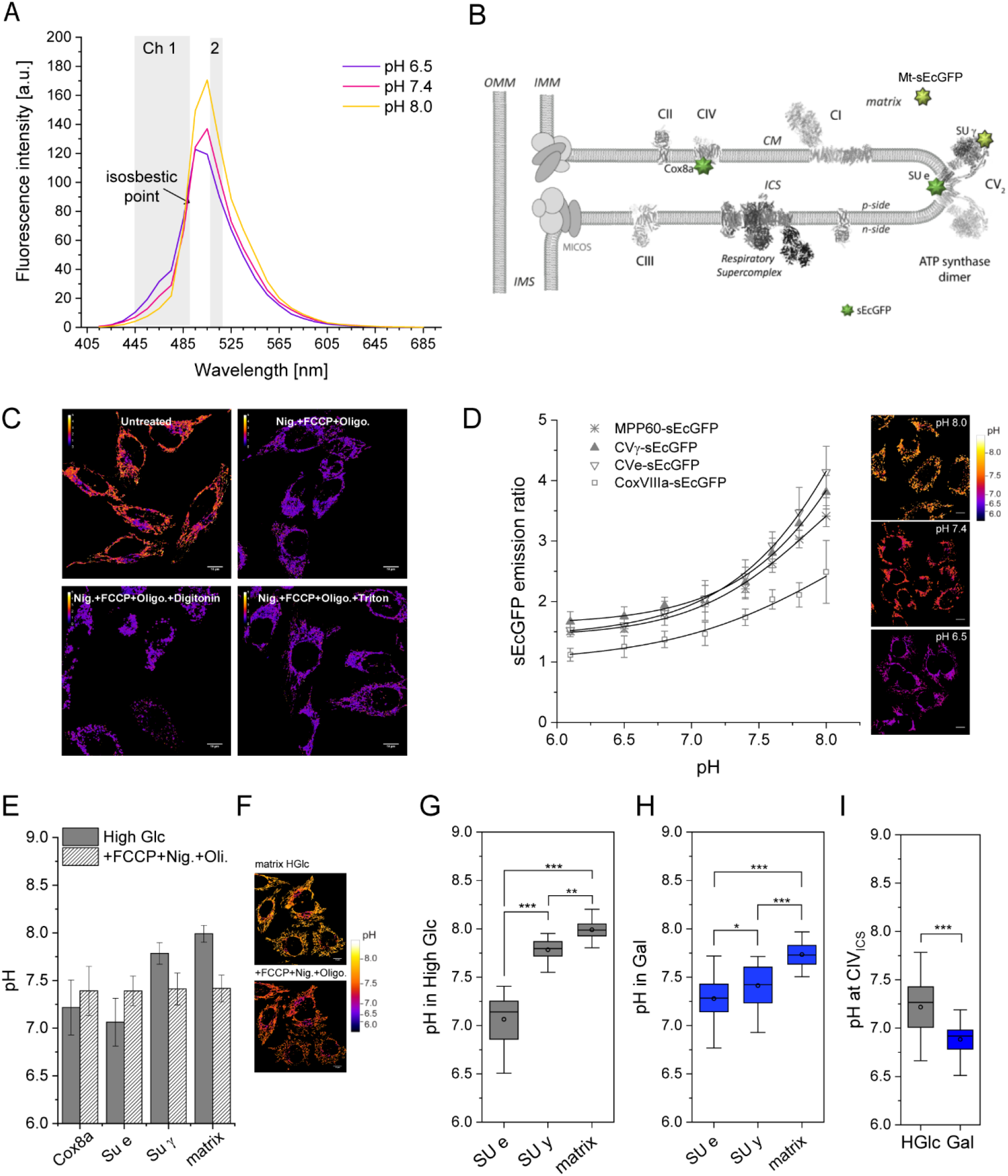
Local pH values in glycolytic and oxidative mitochondria. **(A**) Fluorescence emission spectrum of mt-sEcGFP *in situ* at different pH (λ_ex._ =405 nm, λ_em._ =415-705 nm (λ-emission scan of transfected cells with 10 nm steps using a cLSM microscope). Before and after the isosbestic point (indicated) the pH dependence of the emission is reverse allowing for the read out of pH-dependent fluorescence emission in the two indicated channels and ratio-metric imaging. The λ-range for the first channel is larger than the λ-range for the second channel allowing a better match of the signal intensities of the two different channels. **(B)** Localization of site-specific pH sensors used in this study: fusion of pH-sensitive sEcGFP to subunit e of complex V (SU e-sEcGFP) in the ICS (*p*-side of IMM), to subunit γ of CV (SU γ-sEcGFP) at the *n*-side of the IMM, matrix targeted sEcGFP (mt-sEcGFP) and to subunit CoxVIIIa of complex IV (CoxVIIIa-sEcGFP). For generation of mt-sEcGFP, sEcGFP was fused to the matrix targeting sequence of MPP60. **(C)** Equilibration of intra- and extracellular pH for pH-calibration studies. FCCP (10 µM), nigericin (1 µM) and oligomycin (5 µg/mL) were used to equilibrate intracellular with extracellular pH. Detergents (digitonin and Triton-X100) were tested in addition. Stable transfected cells expressing subunit mt-sEcGFP were exposed to the buffer MES (2-(N-morpholino)ethanesulfonic acid, 20mM) with pH adjusted to pH=6.0. The intracellular pH was measured in the presence and absence of the uncoupler FCCP (10µM), the inhibitors nigericin (1 µM) and oligomycin (5 µg/mL) and the detergents digitonin and Triton x-100, respectively. Ratio images (λ_511_ /λ_464_), false color indicates ratio value. **(D)** Calibration curves for the different sensor-fusion proteins. Ratiometric response of sEcGFP to the local pH. Extracellular buffers with different pH used: MES (2-(N-morpholino)ethanesulfonic acid, 20mM, pH 5.6-6.1), BES (N,N-bis(2-hydroxyethyl)-2-aminoethanesulfonic acid, 20 mM, pH 6.1-7.6) or HEPPSO (4-(2-hydroxyethyl)-piperazine-1-(2-hydroxy)-propanesulfonic acid, 20 mM, pH 7.1-8.0) in 125 mM KCl, 20 mM NaCl, 0.5 mM CaCl_2_, 0.5 mM MgSO_4_ plus FCCP (10 µM), nigericin (1 µM) and oligomycin (5 µg/mL) Each data point represents n=2 independent experiments and mean values ± s.d. of at least 15 cells. **(E)** Local pH values at CoxVIIIa, SU e, SU γ and in the matrix bulk in cells supplied with high glucose (High Glc, 25 mM) before and after the addition of FCCP, nigericin and oligomycin demonstrating pH equilibration with extracellular pH under these conditions. **(F)** Ratio-metric fluorescence images of mt-sEcGFP before and after addition of the pH-equilibration cocktail; pH color coded. **(G)** Mitochondrial pH values at SU e, SU γ and in the matrix bulk in hyper-glycolytic cells (High Glc, 25 mM). (**H**) pH at SU e, SU γ and in the matrix in respiring cells (Gal, 10 mM). **(I)** pH at complex IV in the ICS before and after stimulation of the respiratory chain. Box (75%) and Whiskers Plots. Outliner included. Statistics: One Way ANOVA with post hoc Scheffé test: ***, *p*≤0.001; **, *p*≤0.01; *, *p*≤0.05).

### Inhibition of ATP synthase with oligomycin and IF1 discloses forward and reverse activity

For studying the effect of ATP synthase on local pH, an IF1-KO cell line (Salewskij, Rieger et al., 2019) and a cell line expressing the constitutively active variant IF1-H49K were generated (Garcia-Aguilar & Cuezva, 2018, Schnizer et al., 1996). Levels of IF1 were determined by immuno-detection of the IF1 peptide levels from the different cell lines confirmed successful knockout and expression IF1-H49K, respectively (Supplementary Figure S3A,B). In IF1-H49K expressing cells, the total IF1 level was ∼ 2fold increased compared to HeLa WT, while endogenous IF1 was downregulated. IF1 and IF1-H49K bound to the ATP synthase also under OXPHOS conditions (Supplementary Figure S3C)

Increase of the pH at SU e upon oligomycin addition is a strong evidence of active ATPase. Under conditions when ATP is hydrolyzed and protons are pumped into the ICS, the pH at SU should increase after the addition of oligomycin. In contrast, decrease of the pH at SU e is expected when ATP synthesis activity has been blocked by oligomycin addition. When we applied oligomycin under glycolytic conditions, the pH at SU e increased significantly (from pH=7.06 ± 0.25 to pH=7.32±0.16; *p*=4.9*10^−5^), while it decreased after oligomycin application under respiratory (OXPHOS) conditions (from pH=7.28 ± 0.23 to pH=6.8±0.32; *p*=1.3*10^−13)^). This is in accordance with ATPase activity in high glucose and ATP synthase activity under OXPHOS conditions (Figure 2A). In glycolytic IF1-KO cells, oligomycin had the same effect as in wild type cells, an increase of pH from pH=6.95±0.39 to pH=7.31±0.36 (*p*=2.8 *10^−6^) (Figure 2B). This suggests that the acidic pH before oligomycin addition was due to reverse proton pumping by ATPase activity. Surprisingly, under OXPHOS conditions, we found no difference in local pH at SU e with and without oligomycin in IF1-KO cells. However, the pH at SU e was higher in IF1-KO cells under OXPHOS conditions (IF1-KO: pH=7.39±0.28; wt: pH=7.28±0.24; *p*=0.038) (Figure 2C). The reverse tendency was observed under glycolytic conditions. In wt-cells, the pH was pH=7.06±0.25 and in IF1-KO cells pH=6.95±0.39 (*p*=0.099). This indicates higher ATPase activity in IF1-KO under glycolytic conditions, due to missing inhibition of ATPase function.

**Figure 2.**
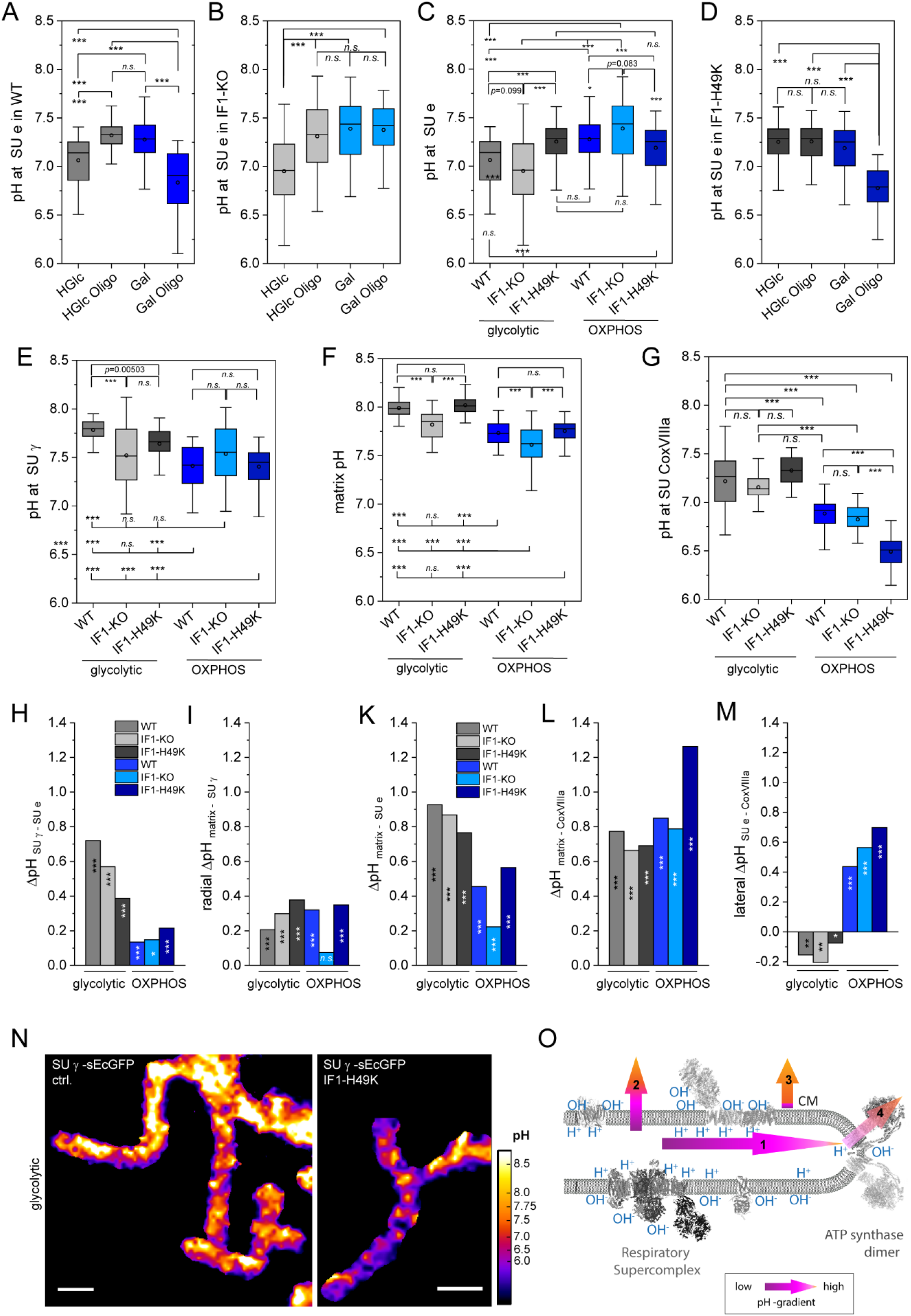
The activity of ATP synthase determines lateral, radial and transmembrane pH gradients. **(A)** Mitochondrial pH at CV SU e measured in HeLa WT in glycolytic (HGlc) and respiring (Gal) conditions. Where indicated, 5 µg/mL oligomycin is added to inhibit CV activity. **(B)** Mitochondrial pH at CV SU e in IF1-KO cells. Conditions: glycolytic (HGlc) and respiring (Gal) with and without oligomycin. **(C)** Mitochondrial pH at CV SU e compared for HeLa WT, IF1-KO and IF1-H49K-HA under glycolytic and respiring conditions. **(D)** Impact of IF1-H49K, the constitutively active IF1 on local pH at SU e. The impact of IF1 is compared between glycolytic and respiring cells. In addition, oligomycin effects are tested. (**E**) Mitochondrial pH at CV SU γ measured in HeLa WT, IF1-KO and IF1-H49K cells under glycolytic and respiring conditions. **(F)** pH in the matrix bulk (MPP60 as mitochondrial targeting sequence) measured in HeLa WT, IF1-KO and IF1-H49K cells under glycolytic and respiring conditions. **(G)** pH at respiratory complex IV at SU CoxVIIIa in the ICS measured in HeLa WT, IF1-KO and IF1-H49K cells under glycolytic and respiring conditions. (**H)** pH differences between SU e and SU γ of ATP synthase SU e is part of F_O_, SU g of F_1_. Triple stars in columns indicate that the difference between the compared pH values constituting the ΔpH was significant; p≤0.001. **(I)** Radial pH gradient between matrix bulk and subunit γ of ATP synthase. (**K**) pH difference between SU e of ATP synthase and matrix bulk. (**L**) Transmembrane pH difference between local pH at CIV SU CoxVIIIa and pH in the matrix bulk. (**M**) Lateral pH gradient between CIV SU CoxVIIIa and CV SU e. Be aware that the pH under glycolytic conditions at SU e was lower than the pH at CIV in the ICS resulting in a “negative” ΔpH. **(N)** pH measurements in mitochondria in live cells expressing sEcGFP as the pH sensor at SU γ. The pH was color-coded according to respective emission ratio. The pH at SU γ in IF1-H49K expressing cells is compared with WT cells. (Scale bar: 1 µm). **(O)** Schematic drawing showing the resolved pH gradients: lateral (arrow 1), transmembrane (arrows 2 and 4) and radial (arrow 3). Statistics: One-way ANOVA with post hoc Schéffe, test for normal distribution. (H-M): One-way ANOVA comparison of sets of pH values at the indicated localizations. Box (75%) and Whisker plots, median (horizontal line in box), mean (box in box). p≤0.001: ***, p≤0.01: **, p≤0.05: *; *n*.*s*.: non-significant (N=3).

### IF1-blocking of ATPase results in increased pH at SU e

Next, we determined local pH at ATP synthase subunit e in cells stably expressing IF1-H49K, a constitutively active IF1 variant(Garcia-Aguilar & Cuezva, 2018, Schnizer et al., 1996), which should inhibit ATPase function of ATP synthase. Blocking of ATPase by IF1-H49K should elevate the pH at SU e, when only ATP synthase is active. Indeed, under glycolytic conditions, the pH at SU e was significantly higher in IF1-H49K cells compared to control cells and IF1-KO cells (Figure 2c), since ATPase was now inhibited and no protons were pumped into the ICS by reverse ATP synthase activity. Oligomycin addition had no further effect on local pH at SU e in IF1-H49K, indicating that previous ATP hydrolysis was already blocked by IF1-H49K (Figure 2D). The pH under OXPHOS conditions was similar as in high glucose (Figure 2C,D). Oligomycin addition resulted in acidification at SU e, since proton were no longer translocated from the ICS to the matrix (Figure 2D). When IF1-KO and IF1-H49K cells were compared, the difference between pH at SU e was significant: The pH value of SU e in IF1-KO cells was pH= 7.39 ± 0.28) and pH= 7.19 ± 0.25 in IF1-H49K expressing cells (*p*< 0.0001) (Figure 2C). This demonstrates that the presence or absence of IF1 expressively affects the pH at SU e of ATP synthase also under OXPHOS conditions. Moreover, a positive effect of IF1-H49K in the presence of endogenous IF1 implicates that endogenous IF1 is not maximally activated at basic pH and when the phosphorylation potential is high.

In order to gain more insights on local pH at ATP synthase in dependence on IF1, we determined pH values at subunit γ of ATP synthase in WT, IF1-KO and IF1-H49K cells. The C-terminus of SU γ fused to sEcGFP extrudes from the F_1_-head of the ATP synthase and the sensor is thus located near at the *n*-side of the IMM. Reverse to the pH at SU e, the pH at CV SU γ in WT and IF1-H49K cells was higher under glycolytic than under respiratory conditions. In IF1-KO cells, the pH was the same under glycolytic and OXPHOS conditions (Figure 2E). Under glycolytic conditions, the presence of IF1-H49K resulted in a pH decrease at SU γ. Under OXPHOS conditions, IF1 had no effect on the local pH at SU γ, the pH was the same in IF1-KO and IF1-H49K overexpressing cells. Since the pH likely differs between membrane surface and matrix bulk(Heberle, Riesle et al., 1994), we also determined the pH values in the matrix bulk in the different cell lines in both metabolic states (glycolytic and OXPHOS). The matrix bulk pH was always lower under OXPHOS than under high glucose conditions. In IF1-KO cells, matrix pH was lowest under glycolytic and OXPHOS conditions (Figure 2F). At the current state, we cannot interpret the drop of pH at the *n*-side and in the matrix in IF1-KO cells without further knowledge. IF1-H49K expression had no influence on the matrix pH in OXPHOS mitochondria; this was expected when only ATP synthase had been active. Under glycolytic conditions no pH difference was found between WT and IF1-H49K expressing cells (Figure 2F). This could be explained by the high buffering capacity of the matrix.

### The local pH at complex IV is not affected by active ATPase

Next, we compared the pH values measured at respiratory complex IV (CIV) in the different IF1 cell lines under glycolytic and OXPHOS conditions. Therefore, sEcGFP was fused to subunit CoxVIIIa at the C-terminus localized at *p*-side (Figure 2G). Under glycolytic conditions, the pH at CoxVIIIa was significantly higher than under OXPHOS condition, since less proton were pumped into the ICS. Higher OXPHOS activity linked with proton pumping resulted in local acid pH values below pH<7. IF1 had no effect on pH values at CIV in glycolytic cells, since neither IF-KO nor IF1-H49K expressing cells exhibited pH differences at CIV. Surprisingly, however, IF1-H49K expressing cells exhibited the most acidic pH at CIV under OXPHOS conditions (pH=6.49 ± 0.17), which was significantly lower than pH=6.88 (± 0.15; *p*<10^−19^) in WT cells (Figure 2G). This suggests either an accelerated cytochrome *c* oxidase proton pumping activity in cells expressing IF1-H49K or a retarded diffusion of protons from the primary proton pump CIV (proton source) to CV (proton sink) (Rieger et al., 2014). Since both, the ultrastructure of the IMM and the spatiotemporal organization of ATP synthase are changed in IF1-H49K expressing cells (Weissert, Rieger et al., 2020), a different kinetic coupling (Toth, Meyrat et al., 2020) is possible.

### ATP synthesis is possible at low Δ*p*

The determined pH values at ATP synthase subunits SU e and SU γ were then used to calculate the corresponding ΔpH. Whether a difference was significant was determined by statistical analysis (Oneway ANOVA test; *p*<0.001: ***). In mitochondria with reverse ATP synthase (=ATPase) activity, the pH difference was highest with ΔpH = 0.75 under glycolytic conditions. IF1-KO and IF1-H49K expressing cells had a lower ΔpH (Figure 2H). In OXPHOS cells, the pH difference was ΔpH ∼ 0.13 under ATP synthesis conditions in WT, IF-KO and IF1-H49K expressing cells. Thus, IF1-H49K reduced the transmembrane ΔpH at ATP synthase under glycolytic conditions, while it did not significantly affect the ΔpH under OPXHOS conditions. Together, these data clearly show that ATPase builds up a local ΔpH at ATP synthase under glycolytic conditions due to reverse proton pumping. In contrast, and quite surprisingly, the local ΔpH and thus the actual Δ*p* was low under steady state OXPHOS conditions at the ATP synthase. This suggests that ATP synthesis is possible at very low ΔpH values: a ΔpH=0.13 means, that the proton concentration at the *p*-side is only 1.1fold higher than at the *n*-side!

### The local pH at SU γ of F_1_F_O_ ATP synthase is lower than in the matrix bulk

Next, we compared the pH between the matrix bulk and the sensor at SU γ (Figure 2E,F, I). Indeed, the pH was lower at SU γ than in the matrix bulk, indicating a radial gradient from the membrane surface towards the bulk. This suggests that proton concentrations near the surface and in the bulk are not in equilibrium, e.g. due to a delay in proton exchange. Such a scenario was proposed before (Heberle et al., 1994, Mulkidjanian, Heberle et al., 2006), and recently intensively reviewed (Lee, 2019). Obviously, diffusion is not sufficient to maintain a homogenous pH in a compartment (Mulkidjanian, 2006) under conditions, when primary and secondary proton pumps are active (Rieger et al., 2014, Strauss, Hofhaus et al., 2008, Toth et al., 2020). This has to be kept in mind, when ΔpH values between the matrix and the IMS/cytosol are calculated: when the matrix bulk pH is taken, the resulting ΔpH is higher (Figure 2H and Figure 2K). Regardless of the reference, though, the ΔpH was lower under OXPHOS conditions compared to glycolytic conditions. Strikingly, in IF-KO cells, the ΔpH was the smallest under OXPHOS conditions pointing to an important role of IF1 in maintaining Δ*p*.

Next, we calculated the ΔpH between CIV and the matrix. Irrespective of the metabolic condition, the calculated ΔpH>0.6. In IF1-H49K expressing cells, the calculated pH difference was ΔpH>1.2 under OXPHOS conditions. This was due to the exceptional low pH at CIV (Figure 2G, K).

### Lateral pH gradients between primary proton pumps and ATP synthase reverse under glycolytic conditions

Finally, we compared pH values at subunit CoxVIIIa at CIV and subunit SU e at ATP synthase. Previously, we and others reported a lateral pH gradient in the IMS (Rieger et al., 2014, Toth et al., 2020). As observed before, the pH difference was ΔpH>0.4 under OXPHOS conditions with a higher proton concentration at CIV. This corresponds > 3fold higher proton concentration at CIV, the primary proton pump (Figure 2M). Under glycolytic conditions, however, the calculated ΔpH_CoxVIIIa_-_SU e_ was negative indicating an inverse pH gradient with a lower pH (higher proton concentration) at SU e of ATP synthase. The inverse lateral pH gradient was also found in IF1-KO and IF1-H49K cells. This can only be explained by the reverse activity of ATP synthase, pumping protons into the IMS while hydrolyzing ATP.

Together, the experimentally obtained pH profiles of the mitochondrial sub-compartments (Figure 2N) revealed that the Δ*p* in mitochondria is not homogenous due to lateral and radial pH. The schematic overview in Figure 2O depicts the measured pH gradients in mitochondria under respiratory conditions. The colored arrows were generated following the ratio-metric pH code used throughout the study. The pH profiles for all conditions are schematically depicted in Supplementary Figure S4.

### IF1 is required to optimize mitochondrial ATP production rates under all conditions

Finally, we correlated mitochondrial and glycolytic ATP synthesis rates and mitochondrial ATP level in living cells with the ΔpH data. To obtain ATP synthesis rates, OCR and ECAR were recorded before and after serial addition of mitochondrial inhibitors (oligomycin and rotenone/antimycin A) and pathway-specific ATP production rates were calculated (Mookerjee, Gerencser et al., 2017). Under glycolytic conditions, HeLa WT cells showed low mitochondrial ATP production rates (<400 pmol/min*30 K cells) but high cytosolic ATP production rates (>600 pmol/min*30 K cells)(Figure 3A,B). To consider possible changes of mitochondria size and number, the mitochondrial area per cells was determined from MitoTracker™Green stained mitochondria (Supplementary Figure S5). OCRs were then normalized on the mitochondrial mass. Under OXPHOS conditions ATP synthesis rates increased >2fold, while cytosolic ATP synthesis rates decreased significantly. The increase in ATP synthesis rates under OXPHOS conditions was not observed in IF-KO cells, though, emphasizing the role of IF1 to block ATP hydrolysis under OXPHOS conditions due to low ΔpH/Δ*p*. Also, ATP levels were lower in IF1-KO cells under OXPHOS conditions (Figure 3C).

**Figure 3.**
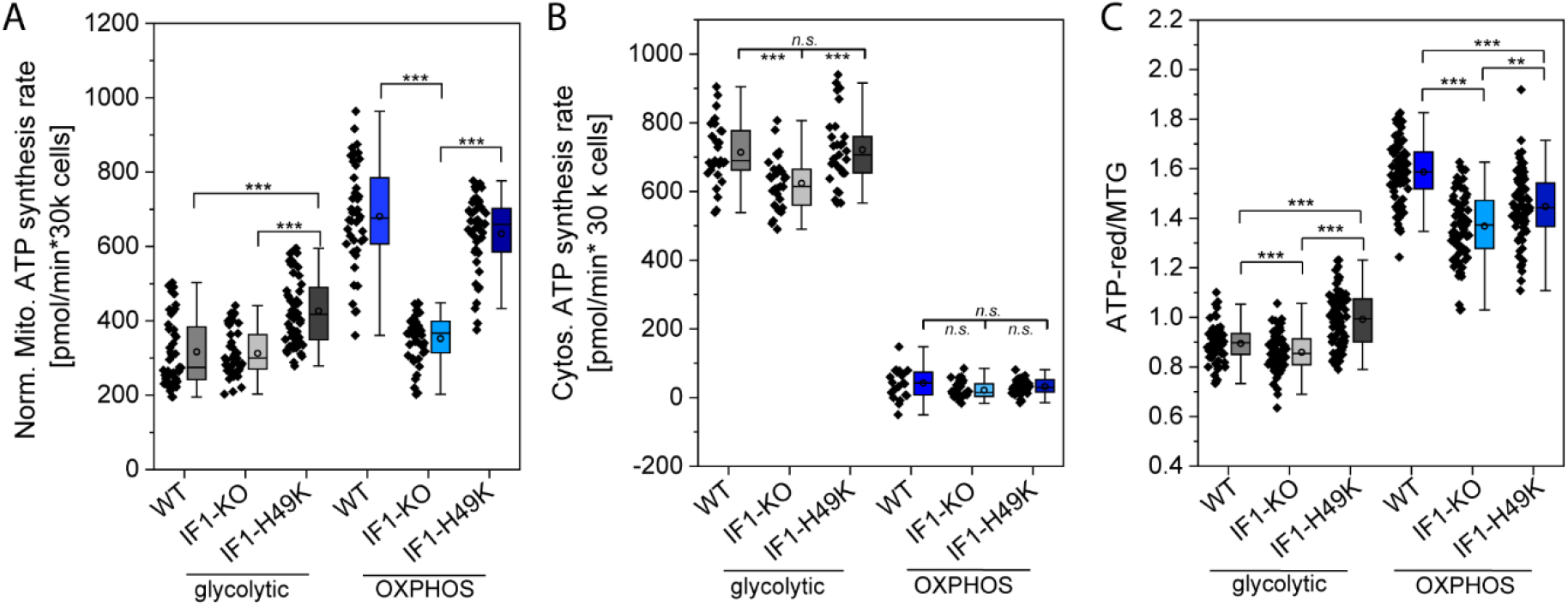
IF1 is beneficial for ATP production even under OXPHOS conditions. Mitochondrial ATP production rates in the presence and absence of IF1. Oxygen consumptions rates (OCR) and extracellular acidification rates (ECAR) were determined with an automatic flux analyzer (Seahorse/Agilent). After monitoring basal respiration, Oligomycin (1 µM), and rotenone plus antimycin A (Rot; 0.5 µM; AA, 0.5 µM) were added sequentially to determine glycolytic and mitochondrial ATP production rates. **(B)** Cytosolic ATP production rates. Each data point represents the mean ATP production rate of one well (N=3). **(C)** Determination of mitochondrial ATP levels with the fluorogenic dye ATP-red™ (5 µM, prestaining for 15 min). To normalize on mitochondrial mass, mitochondria were stained with MitoTracker™ Green (MTG, 100 nM for 30 min). Statistics: One Way ANOVA with post hoc Scheffé test; ***, p≤0.001; **, p≤0.01; *, p≤0.05; n.s.: non-significant.

Under glycolytic, but not OXPHOS conditions, IF1-H49K had a positive effect on ATP production rates. We conclude that the endogenous IF1 was not fully active under glycolytic conditions, likely because the matrix pH was basic (pH>7.5) and IF1 is activated at pH<7.5 (Boreikaite et al., 2019, Zanotti, Gnoni et al., 2009). The pH-independently active IF1 (IF1-H49K) obviously blocked ATPase more efficiently. Finally, ATP levels were determined with the fluorogenic BioTracker™ ATP-Red™dye (Millipore) which locates to mitochondria. ATP-Red fluorescence was normalized on the mitochondrial mass (MitoTracker™Green signal). Generally, mitochondria with high respiration rates (OXPHOS conditions) had a significant higher ATP content than mitochondria with lower respiration (glycolytic conditions), irrespective of IF1 levels (Figure 3C). IF1-H49K expressing glycolytic cells had more mitochondrial ATP than WT cells in accordance with higher mitochondrial ATP synthesis rates and inhibition of ATP hydrolysis. IF1-KO cells had lower ATP levels than WT cells which can be explained by higher ATP hydrolysis rates and/or export of ATP to supplement the cytosol with ATP as suggested above. Thus, these data suggest that in WT HeLa cells a subpopulation of ATP synthase is in the reverse mode. Under OXPHOS conditions (Salewskij et al., 2019, Weissert et al., 2020), IF1 is required to block this. Unexpectedly, also IF1-H49K expressing cells had lower mitochondrial ATP levels than WT cells (but higher levels than IF1-KO cells). Whether this is due to a hampered coupling of ATP synthase and respiration could be tested by determining P/O values. A decreased coupling could be due to the recently observed altered spatiotemporal organization of ATP synthase in IF1-H49K expressing cells (Weissert et al., 2020).

Clearly, this data demonstrates the importance of IF1 to block futile ATP hydrolysis under OXPHOS conditions.

## CONCLUSION

ATP synthase activity substantially determines the local Δ*p* in mammalian cells as local pH measurement revealed. Moreover, the Δ*p* at ATP synthase is almost negligible under OXPHOS conditions. To prevent futile ATP synthesis/hydrolysis cycles by reverse ATP synthase activity and ATP hydrolysis, IF1 is essential even and especially under OXPHOS conditions. In addition, pH profiling of mitochondrial sub-compartments disclosed that the pH is neither laterally nor radially balanced in mitochondria in living cells. This is possible due to the physical separation of primary proton pumps and ATP synthase. Obviously, diffusion of protons is not sufficient to ensure proton equilibration under steady state conditions. In order to understand the proton motive force, this has to be taken into consideration.

## AUTHOR CONTRIBUTIONS

R., T.A. and K.B.B. designed the experiments, B.R. performed pH, ATP and respiration experiments, T.A conducted protein gels and immune-blotting, J.V. was involved in pH measurements and B.R. and K.B.B. prepared figures. K.B.B. and B.R. wrote the main manuscript text.

## DATA AVAILABILITY STATEMENT

The data generated and analyzed in the current study are available from the corresponding authors on request.

## COMPETING FINANCIAL INTEREST

The authors declare no competing financial interests.

## ACKNOWLEDGMENTS

The study was supported by grants from the DFG (Bu2288/1-2), (Bu2288/1-3) and the CRC SFB944 (INST 190/1672). We thank Wladislaw Kohl and Patrick Duwe for technical assistance and Giulia Nebel for important preliminary studies. We also would like to thank Michael Hippler and Armen Mulkidjanian for critically reading the manuscript.

## Notes

### Competing Interest Statement

The authors have declared no competing interest.

### Summary of Updates

Title was changed

