## Supplemental Figures and Material for "Mitochondrial F_1_F_O_ ATP synthase determines the local proton motive force in cristae tips"

### 1 MATERIAL AND METHODS

#### Cell culture

HeLa cells were purchased from the Leibniz Institute DSMZ-German Collection of Microorganisms and Cell Cultures). The HeLa cells (wildtype, IF1-KO and IF1-H49K-HA) were cultivated in XF Base Medium Minimal DMEM (Agilent) supplemented with 10 % FBS (Fetal bovine serum), 1 % HEPES, 1 % NEAA (non-essential amino acids), 2% $NaHCO_3$ and either 10 mM D-Galactose (plus 4 mM Alanyl-L-Gutamine) or 25 mM D-Glucose (plus 2 mM Alanyl-L-Gutamine) at 37 °C and 5% $CO_2$. When confluency was reached, cells were split using 1 mL Trypsin/EDTA for 5 min at 37 °C (and 5% $CO_2$).

#### Generation of IF1-KO and IF1-H49K-HA cell lines

The IF1-knockout cells (IF1-KO) were manufactured using the CRISPR/Cas technique. The first step was to find the appropriate targeting sequence for crRNA in the vicinity of the START codon. The crRNA was cloned into the pSpCas9(BB)-2A-GFP (PX458) vector (Addgene). Thus, the resulting vector contained both sgRNA and Cas9 nuclease. To enable selection with Puromycin, the *pac* gene was inserted at the cutting site. Therefore, a construct was cloned with homologous arms to the right and left of the place where Cas9 is to be cut with the pac-gen in between. HeLa cells were co-transfected with both constructs and transformed cells were selected by adding 0.5 µg/mL Puromycin. Positive clones were isolated and the IF1 knockdown tested by qPCR and Western. For cloning of IF1-H49K-HA, the forward primer GAAGAAGAAATCGTTCATCATAAG with an EcoRV restriction site and the reverse primer CTTGTGTTTTTTCAAAGCTGCCAGTTGTTC with an EcoRI restriction site was used. The template was integrated into the pSems26 vector after cutting out the sequence encoding for CoxVIIIa-snap-tag purchased from NEB Biosciences™ (formerly Covalys™). The plasmid backbone of pSems26 encodes for ampicillin resistance and has a CMV promoter. For generation of a stable cell line expressing IF1-H49K-HA, transfected HeLa cells were selected for stable neomycin resistance by growth in the presence of 0.8 mg/mL G418 (Calbiochem 345810). Positive clones were tested by Western.

#### Real-Time ATP Rate Assay via oxygen consumption measurements

Oxygen consumption rates (OCR) and extracellular acidification rates (ECAR) of intact HeLa cells were recorded with the Seahorse XFe96 Extracellular Flux Analyzer (Agilent Technologies). 30.000 cells were sown per well of a 96-well XF cell culture microplate 24 h before the experiment. 60 min before the Agilent Seahorse XF Real-Time ATP Rate Assay, cells were washed with XF DMEM Medium pH 7.4 (with 5 mM HEPES, without phenol red, glucose, pyruvate, and L-glutamine, 103575-100 from Agilent/Seahorse Technologies) with supplements (1 mM pyruvate, 2 mM L-glutamine and 5.6 or 25 mM D-glucose, respectively, 10 mM D-galactose – adjusted to pH 7.4), added to fresh XF assay medium and incubated at 37°C before loading into the XFe Analyzer. Supplements were from Roth. Injections were as follows: *1*. 1 µM oligomycin, *2*. 0.5 µM rotenone/0.5 µM antimycin. Seahorse XF technology measures the flux of both $H^+$ production (ECAR) and $O_2$ consumption (OCR), simultaneously. By obtaining OCR and ECAR data under basal conditions and after serial addition of mitochondrial inhibitors (oligomycin and rotenone/antimycin A), total cellular ATP production rates and pathway-specific mitochondrial ATP and glycolytic ATP production rates can be measured (Mookerjee et al., 2017). The series of calculations used to transform the OCR and ECAR data to ATP production rates is performed using the Seahorse XF Real-Time ATP Rate Assay Report Generator from Agilent. Briefly, glycolytic ATP production rate (pmol ATP/min) = glycolytic proton efflux rate (pmol $H^+$/min) is associated with the conversion of glucose to lactate in the glycolytic pathway. Mitochondrial ATP production rate is associated with oxidative phosphorylation in the mitochondria and defined as: $OCR_{ATP}$ (pmol $O_2$/min) = OCR (pmol $O_2$/min) - $OCR_{Oligo}$ (pmol $O_2$/min).

### SDS-PAGE and Western blotting

For SDS-PAGE cell lysates from samples of confluent T-25 flasks (1 flask would last for 10-15 gels) were heated for 5 min (95°C) and then separated on 12% Tricine-SDS-PAGEs and transferred to PVDF membranes. Membranes were blocked with 10% nonfat dry milk in TBS-T (200 mM Tris, 1.37 M NaCl, + 0.1% Tween20). IF1 was detected with anti-ATPIF1 antibody (CD6P1Q) purchased from Cell Signaling (#13268), VDAC was detected with an anti-VDAC antibody (Cell signaling #4661).

### Fluorescence microscopy

Fluorescence imaging was carried out with confocal laser scanning microscope (Leica TCS SP8 SMD) equipped with a 63× water objective (N.A. 1.2) and a tunable white light laser. HyD's with GaASP photocathodes were used as detectors. Measurements were performed in a humidified chamber at 37 °C and 5% $CO_2$.

### Local pH measurements

For *in situ* pH determination, HeLa cells transfected with sEcGFP (Miesenbock et al., 1998) (or superecliptic pHluorin = pHl, derivate of green fluorescent protein) fusion constructs (calcium phosphate method) were used. sEcGFP can be used as a ratio metric pH sensor (Rieger et al., 2014), when emission at 470 nm and 510 nm are recorded (Gao et al., 2004). The original template for sEcGFP was a gift of Prof. Jürgen Klingauf. The local pH sensors were generated by genetically fusing sEcGFP to subunits of CV, CIV and targeting it to the matrix, respectively (Figure 1D). In detail, sEcGFP was fused to subunit $\gamma$ (SU $\gamma$) of the $F_1$ subcomplex of CV at the *n*-side of the IMM, to subunit e (SU e) of $F_O$ subcomplex of CV at the *p*-side of the cristae membrane and to subunit CoxVIIIa of CIV at the *p*-side of the IMM (Rosselin et al., 2017). To generate CoxVIIIa-sEcGFP, the sEcGFP template was integrated into the pSems26 Cox8a-snap vector purchased from NEB Biosciences™ (formerly Covalys™) after cutting out the sequence encoding for the snap-tag. The plasmid backbone of pSems26 encodes for ampicillin resistance for bacterial selection and contained the neomycin resistance for selection of stable clones. The promoter was from CMV. To generate SU e-sEcGFP and SU $\gamma$-sEcGFP, respectively, CoxVIIIa was cut out and the coding sequences for SU e and SU $\gamma$, respectively inserted at the N-terminus of sEcGFP. The correct localizations of the fusion constructs have been confirmed by Immuno-EM in a different study (Rosselin et al., 2017). An additional pH sensor was generated for the matrix. The mitochondrial targeting sequence of the mitochondrial processing peptidase MPP was fused to sEcGFP to obtain a soluble matrix pH sensor (mt-sEcGFP). Stable cell lines were generated by addition of 0.5 µg/mL Puromycin or 0.8 mg/mL G418 and selection and testing of positive clones. CoxVIIIa-sEcGFP and CV Su e-sEcGFP were used to determine ICS pH and CV, SU $\gamma$-sEcGFP and mt-sEcGFP to determine matrix pH (Rieger et al., 2014; Sohnel et al., 2016). Local pH was recorded *in situ* by confocal fluorescence emission ratio imaging by an inverse confocal laser scanning microscope (Leica TCS SP8 SMD). All measurements were performed at 37°C in fresh medium or buffer. Ratio images were obtained after excitation with a single wavelength ($\lambda_{ex.}$=405 nm) by recording in two wavelength regimes ($\lambda_{em.}$ = 440–490 nm and 505–517 nm). For pH calibration, cells were perfused for 3–5 min with PBS (PAA, pH 7.0–7.5 with $CaCl_2$ and $MgCl_2$). PBS was exchanged for MES (25 mM, pH 6.1), BES (25 mM, pH 6.5, 7.1 and 7.4) or HEPPSO (25 mM, pH 7.8, 8.0 and 8.2) buffer adjusted to the desired pH by 1M HCl or NaOH and supplemented with the following compounds: 125 mM KCl, 20 mM NaCl, 0.5 mM CaCl2, 0.5 mM $MgSO_4$, 10 µM FCCP, 1 µM nigericin and 5 µg/mL oligomycin. FCCP, nigericin and oligomycin were purchased from Enzo Life. MES was purchased from Biomol and BES and HEPPSO from Sigma. Other common chemicals were sourced from Roth. Images were taken after 10-30 min of incubation.

### Mitochondrial ATP via fluorescence imaging of ATP-red

The BioTracker™ ATP-red dye (Millipore) is a fluorogenic indicator for ATP in mitochondria (Wang et al., 2016). To load the dye, the cells were incubated in medium with 5 µM ATP-red for 15 min at 37 °C (5% CO2). Before measurement, the cells were washed twice with medium before fresh medium was added. The fluorescence was recorded with a cLSM (Leica SP8, 60x water objective) at 37 °C and 5% $CO_2$. ATP-red was excited at 561 nm (white light laser) and the emission from Z-stacks (3 slices, step size 360 nm) was recorded between 580 and 650 nm. The subtracted background images for intensity analysis were created with ImageJ® software (NIH Image). The filter mask Otsu was used as mask for the cells and the background was set to NaN.

**Statistics**. Statistical analysis was performed using OriginPro version 9.6, 2019 (OriginLab Cooperation, Northhampton, MA). After testing for normal distribution (Kolmogorov-Smirnov), Oneway ANOVA test with post-hoc Schéffe test was selected for comparative measurements. At least 3 independent measurements on different days with different cell culture samples were the basis.

### pH calibration

sEcGFP has two emission peaks ($\lambda_{em1}$=511 nm and $\lambda_{em2}$= 464 nm, $\lambda_{exc}$=405 nm) that reversely respond to pH changes. The emission spectrum obtained by a fluorescence scan (10 nm steps) of cells stably expressing matrix-targeted sEcGFP shows the existence of an isosbestic point at 484 nm Thus, the fluorescence intensity below and after 484 nm shows an opposite dependency on pH and sEcGFP can be used as ratio-metric pH probe. The fluorescence emission ratio of $\lambda_{em1}/\lambda_{em2}$ reports pH changes[14]. The ratio of Em.511/Em.464 was plotted in dependence on the pH. For all constructs, the $\lambda_{em1}/\lambda_{em2}$ value increased. The dependency was fitted with a Boltzmann fit to obtain the respective conversion from emission ratio to pH.

### Immunoblotting

Briefly, cell lysates from samples of confluent T-25 flasks (1 flask would last for 10-15 gels) were heated for 5 min (95°C). 20 µg of denatured proteins were then resolved by SDS-PAGE using self-casted 4- 12% Tricine-Bis-Acrylmaide gels (BioRad). Gel contents were electrotransferred to PVDF membranes (Millipore) using a semidry TransferApparatus (Trans-Blot SD BioRad). Equal loading was evaluated by Ponceau S staining. Membranes were blocked with 10% nonfat dry milk in TBS-T (200 mM Tris, 1.37 M NaCl, + 0.1% Tween20) for 1 h. Membranes were incubated with primary antibodies overnight at 4°C. IF1 was detected with anti-ATPIF1 antibody (CD6P1Q) purchased from Cell Signaling (#13268), VDAC was detected with an anti-VDAC antibody (Cell signaling #4661). After washes in TBS-T, detection was performed with HRP-conjugated secondary antibodies (1:2000). Membranes were washed in TBST and developed using standard chemiluminescence with ECL (SuperSignal WestPico Thermofisher) and imaged by ChemiDoc BioRaD.

### 2 SUPPLEMENTARY TABLES

**Table 1: Boltzmann fitting for the pH sensor-proteins**

| y=A2+(A1-A2)/(1+exp((x-x0)/dx))<br>Modell Boltzmann | | | | |
|---|---|---|---|---|
| | matrix | SU [illegible] | SU e | CoxVIIIa |
| A1 | 1.4212<br>± 0.0779 | 1.6212<br>± 0.1056 | 1.4149<br>± 0.0529 | 0.9945<br>± 0.1876 |
| A2 | 5.1628<br>± 1.9465 | 10.26836<br>± 13.6406 | 12.7694<br>± 9.2931 | 4.30476<br>± 5.0804 |
| X0 | 7.9397<br>± 0.4560 | 8.5303<br>± 1.2105 | 8.6165<br>± 0.6513 | 8.1848<br>± 1.7281 |
| dx | 0.4656<br>± 0.1398 | 0.4976<br>± 0.1820 | 0.5379<br>± 0.0831 | 0.6561<br>± 0.4623 |
| Chi-Quadr. Red. | 0.3736 | 0.1196 | 0.0414 | 0.1369 |
| $R^2$ (COD) | 0.994 | 0.994 | 0.998 | 0.987 |

### 3 SUPPLEMENTARY FIGURES

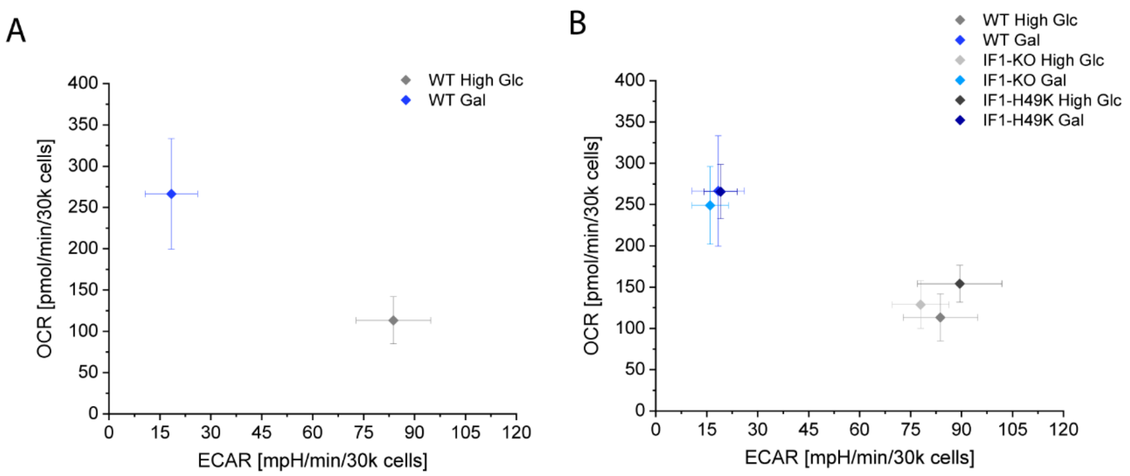


**Supplementary Figure S1. Metabolic setting in dependence on sugar supply and IF1-levels.**

(A) Determination of OXPHOS and glycolytic activity by monitoring oxygen consumption rates (OCR) and extracellular acidification rates (ECAR). OCR and ECAR were determined with an automatic flux analyzer (96XF, Seahorse/Agilent). ~30,000 cells per well were seeded the day before measurement. Before measurements, cells were supplied with glucose (25 mM, HGlc) or galactose (10 mM, Gal) for 3 weeks.

(B) Metabolic profile in dependence on IF1 levels under glycolytic and OXPHOS conditions.

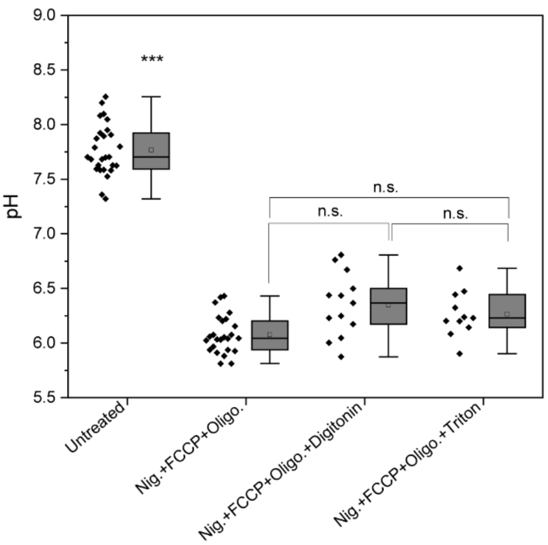


**Supplementary Figure S2. pH equilibration between mitochondria and extracellular pH does not require detergents.**

Cells expressing mt-sEcGFP were imaged in the absence and presence of a cocktail of different inhibitors (oligomycin: 5 µg/mL) and uncouplers (FCCP: 10 µM and nigericin: 1 µM) with and without detergents (digitonin: 0.006 w/v or Triton-X100: 0.002 w/v). The pH of the external medium was pH 6.1. One-way ANOVA comparison of pH values in the matrix after treatment of cells with the cocktails for pH equilibration. Box (75%) and Whisker plots, median (horizontal line in box), mean (box in box). $p \leq 0.001$: ***, *n.s.*: non-significant (N=3).

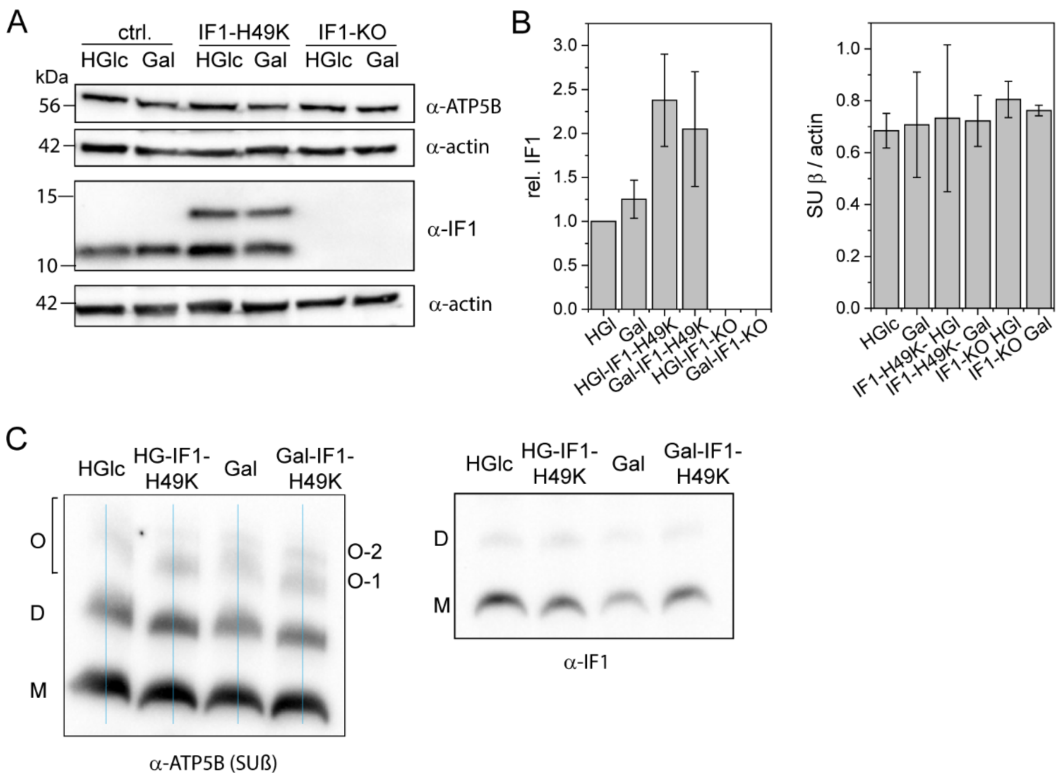


**Supplementary Figure S3. IF1-levels in cell lines used in this study.**

(A) Immuno-blotting of IF1 to determine IF1 levels in HeLa WT, in IF1-KO and in IF1-H49K-HA expressing cells. VDAC was used as loading control.

(B) Relative IF1 levels of cells used in this study.

(C) IF1 binds to ATP synthase under different metabolic conditions. BN-PAGE, immunoblotting with α-(ATPB) and anti-IF1, respectively. O: oligomers, D: dimers; M: monomers.

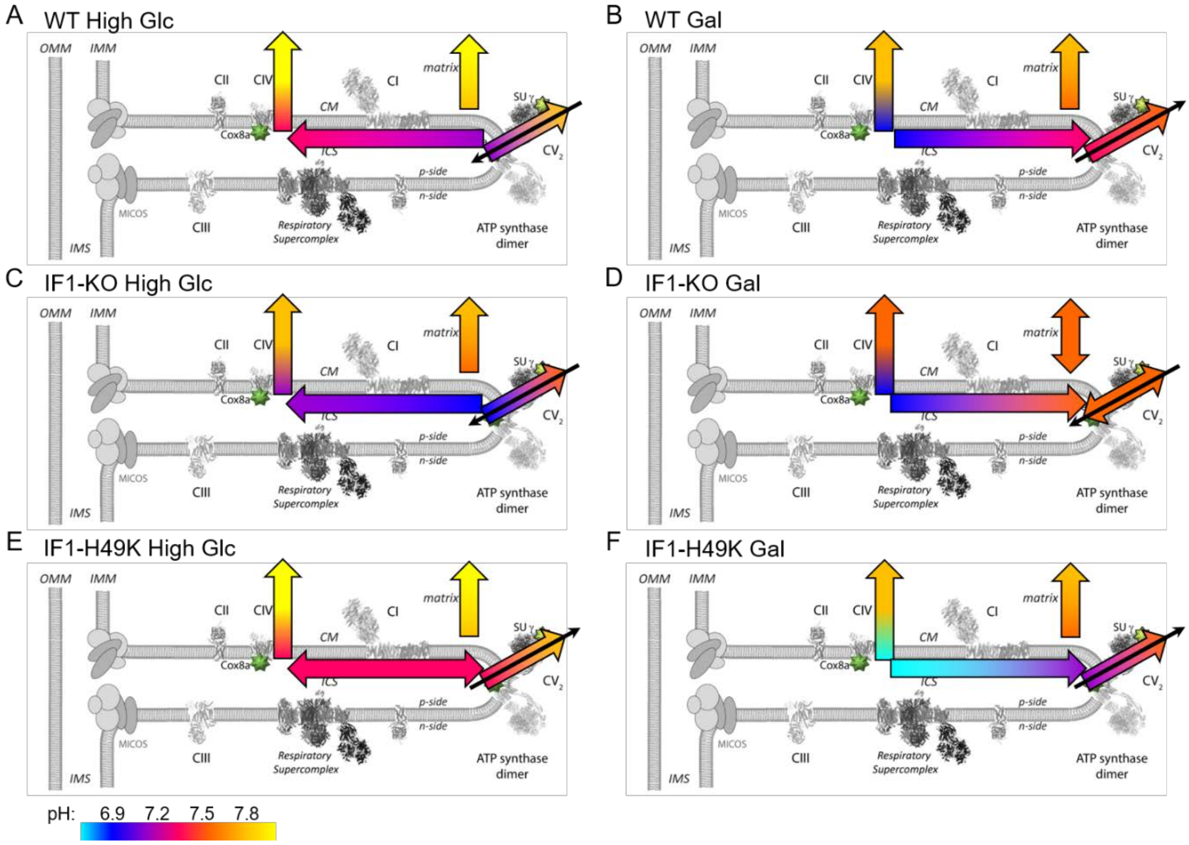


**Supplementary Figure S4. Influence of IF1 on the transmembrane, radial and lateral pH gradients in mitochondria under glycolytic and oxidative conditions.**

(A) Local protonic energy coupling in glycolytic cells.
(B) Local protonic energy coupling in respiratory cells.
(C) Local protonic energy coupling in glycolytic cells without the ATP synthase inhibitor IF1.
(D) Local protonic energy coupling in respiratory cells without the ATP synthase inhibitor IF1.
(E) Local protonic energy coupling in glycolytic cells with constitutively active IF1.
(F) Local protonic energy coupling in respiratory cells with constitutively active IF1.

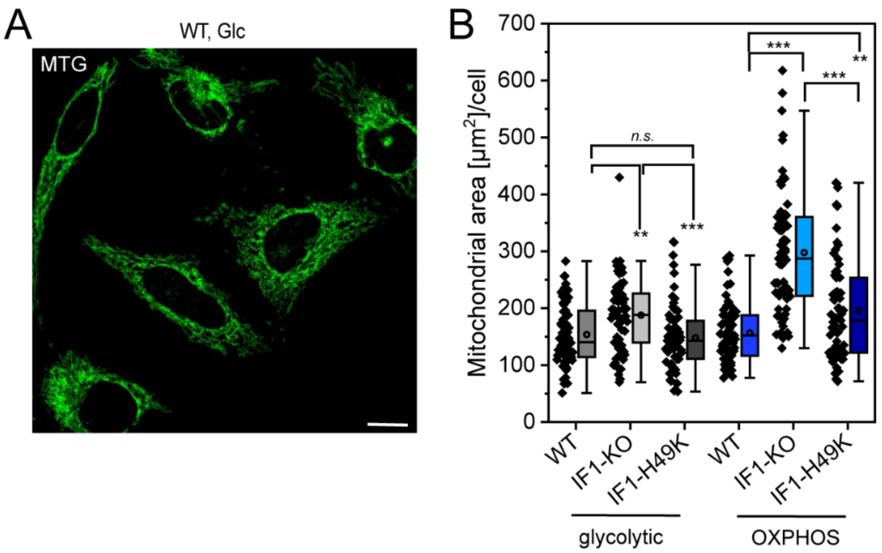


**Supplementary Figure S5. Mitochondrial area per cell under different conditions.**

(A) Determination of mitochondrial area. Cells were stained with MitoTracker™Green (MTG, 100 nM for 30 min) to visualize mitochondria. Mitochondrial area was determined by integrating grey values deriving from MTG from cell.

(B) Mean mitochondrial area per cell for different cells lines and metabolic conditions. Scale bar: 10 µM (a). Statistics: One Way ANOVA with post hoc Scheffé test; ***, p≤0.001; **, p≤0.01; *, p≤0.05; n.s.: non-significant. Box covers 25-75 perc. Extremes are shown, rectangle indicates mean, horizontal line indicates median.
